# Paternal Indifference and neglect in early life and Creativity: Exploring the Moderating Role of *TPH1* genotype and Offspring’s Gender

**DOI:** 10.1101/728600

**Authors:** Qi Yu, Si Si, Shun Zhang, Jinghuan Zhang

**Affiliations:** Department of Psychology, Shandong Normal University, Jinan, China

**Keywords:** Creativity, Paternal indifference & neglect in early life, *TPH1*, Gene-environment interaction

## Abstract

For further understanding the joint contribution of environment, heredity and gender to creativity, the present research examined the prospective impact of paternal indifference & neglect in early life, *TPH1* rs623580, offspring’s gender, and the interaction effects thereof on creativity in five hundred and thirty-nine unrelated healthy Chinese undergraduate students. Paternal indifference & neglect in early life was assessed on the Parental Bonding Instrument (PBI) and creativity on the Runco Creativity Assessment Battery (rCAB). Results showed significant paternal indifference & neglect × *TPH1* genotype and *TPH1* genotype × offspring’s gender interaction effects when predicting creativity. Specifically, paternal indifference & neglect in early life negatively predicted creativity in youth when individuals carry A allele of *TPH1* (rs623580). In addition, male individuals who carry A allele were linked with lower level of flexibility compared to TT homozygote individuals. No significant three-way interaction was found. Findings from the current study suggested that the A allele of *TPH1* (rs623580) might be a risk allele for creativity, and the long-term negative influence of paternal indifference & neglect in early life on individuals’ creativity in youth depending on *TPH1* genotype.

## INTRODUCTION

Creativity is defined as the capacity for producing something that is both novel and useful [1–3].There is a consensus in the field that creativity involves in the improvement of technology, science, art, philosophy, or even all walks of life [4].Previous studies indicated that creativity was the major driving forces behind the progress of civilization [5, 6].

Because of the central role creativity plays, there has always been a great interest for psychologist on how biological and environmental factors foster or inhibit creativity [7, 8]. For the biological factors, recent advances in molecular genetics have permitted psychologists to explore the underlying genetic basis of creativity, and several genes (e.g. *THP1, TPH2*) were revealed to associate with creativity [9–11].

For the environmental factors, parenting is one of the most frequently investigated due to its crucial role in creativity [12, 13]. However, results from twin and adoption studies have indicated that creativity cannot be explained exactly by either gene or environment [14, 15]. A growing evidence highlighted the importance of Gene × Environment (G × E) interactions, in which the relationship between environmental factors (e.g. parenting) and child outcomes (e.g. antisocial behaviors, cognitive abilities, social function, wellbeing) might be moderated by genetic factors [16, 17]. Therefore, the primary purpose of present study was to explore the interaction effect of genetic and environmental factors on creativity.

Besides, previous studies indicated that gender difference might be attributed to the interaction effect of genetic and environmental factors on creativity [18]. Therefore, gender of offspring was another variable recruited in this study, exploring the possiple Gene × Environment × Gender (G × E × G) interaction effect when predicting creativity.

### Parental indifference & neglect and creativity

One vital factor that has long been recognized to influence creativity is the early life family environment, among which parenting have received the most attention [19–21]. Parental indifference & neglect is a significant risk factor for children across their psychological and behavioral development and is usually linked with several serious aftermath that reach into adulthood [22–24], including psychological maladjustment, internalizing/externalizing behaviors, and negative personality dispositions of children [22, 25, 26].

According to Parental Acceptance-rejection Theory (PART), parental indifference refers to a mood state of parents distinguished by a lack of care, concern and interest of their children; while parental neglect refers to a behavioral response that parents fail to attend the physical, psychological, and social needs of their children appropriately [25, 27]. Although the connection between indifference and neglect is not extremely direct, such as parents may neglect or be perceived to neglect their children for many reasons which are not driven by indifference, both indifferent and neglecting parents remain unavailable and unresponsive to their children’s need, consistently [6]. While most of the attention in the field of parental indifference & neglect has been directed toward negative outcomes, evidences provided by recent empirical studies have indicated that parental indifference & neglect in early life negatively predicted positive outcomes, such as cognition and intelligence [28–31]. Using the Audio-Computer Assisted Self Report Interview (ACASI), one study investigated the relation between multidimensional neglect and cognition, the result showed that children suffering neglect had lower overall coginitive performance in comparison with normative data [30]. Coincidentally, using the Wechsler Intelligence Scale for Children -Revised, Split-Half Short Form (WISC-R:SH), Kaufman et al. (1994) reported a direct relation of neglect to intelligence quotient (IQ), with children who experiencing the most severe parental neglect having the lowest performance in IQ scale [31]. A further study demonstrated that the neglected children showed lower general intelligence and poorer executive decision than the controls [28]. Creativity and divergent thinking are deemed to be facets of intelligence in some intelligence models [9, 32, 33]. So based on the notion, parental indifference & neglect in early life might play a negative prospective role in creativity in youth.

However, the existing parenting research has documented that parental indifference & neglect in early life is not always deleterious, especially in creativity research. Previous studies provided evidence that parental indifference & neglect may positively relate to child’s creativity. Albert (1992) reported that many genius and great eminences were suffered from parental indifference & rejection and poverty in early family environment [34]. Similarly, a longitudinal study, which aimed to reveal the association between parent-child relationships and creative personality traits, suggested that individuals with creative personality traits, such as self-sufficient, reserved, serious, adventurous, and sensitivity, were inclined to report their parents expressed more neglect and reject while they were growing up [35]. Inconsistent findings suggest that the relation between parental indifference & neglect in early life and child developmental outcomes may be moderated by additional variables.

One possible explanation is that the influence of parental indifference & neglect to children may be differ between mother and father. However, much studies in this research area were conducted with both mother and father [25, 36], few studies examined specially fathers’ indifference & neglect and its influence on child developmental outcomes [37, 38]. An ever-expanding line of research has indicated that fathers played an important role in children’s psychological and behavioral development, including academic achievement, cognitive development, behavioral or emotional regulation and so forth [39, 40]. Thus, the present study was designed to investigate the particular relation of paternal indifference & neglect in early life to creativity in youth.

Moreover, previous studies indicated that if father was unavailable, then boys had a greater likelihood of engaging in negative outcomes [41, 42]. Given that father is the most significant model for boys’ identification [43], it is logical to infer that the role of paternal indifference & neglect in offspring developmental outcomes may be different for boys and girls. Therefore, it is possible that offspring’s gender may moderate the relationships between paternal indifference & neglect in early life and creativity in youth.

### *TPH1* rs623580 and creativity

Studies utilizing behavior genetic research designs demonstrated both genetic and environmental factors have influence on individual’s creativity [44]. Recent advances in molecular genetic studies have permitted direct exploring the underlying mechanism of the G × E interaction via identifying specific genes or locus associated with creativity. Empirical research showed a genetic variant in the dopamine D2 receptor gene *(DRD2)*, rs1799732 polymorphisms, moderated the relation between authoritarian parenting and creativity [45]. Therefore, we postulated in this line that the relation between paternal indifference & neglect in early life and creativity in youth may be moderated by genetic variants.

Besides *DRD2*, several lines of research indicated the TPH1genotypes involve in creativity. Using inventiveness battery of the Berlin Intelligence Structure Test (BIS), Reuter et al. (2006) reported that *TPH1* rs1799913 (A779C) polymorphism was significantly associated with creativity. Similar findings, using Divergent Thinking Test (DT Test), indicated that *TPH1* rs1799913 polymorphism was significantly associated with ideational fluency [10]. To further elucidate the role of *TPH1* in creativity, by including both related functional SNPs and tag SNPs, a recent study comprehensively explored the correlation between *TPH1* genetic variants and creative potential measured by DT Test [11]. Although failed to replicate the correlation of *TPH1* rs1799913 and creativity, the results suggested a new *TPH1* genetic variate, rs623580 (T3804A), associated with both verbal and figural fluency.

*TPH1* rs623580 located in the exon 1c & intron1 within the 5’-UTR of the *TPH1* gene at human chromosome 11 [46]. *TPH1* is the rate limiting enzyme in the biosynthesis pathway of the neurotransmitter 5-hydroxytryptamine (5-HT, Serotonin) and therefore a critical step in 5-HT functioning [47]. *TPH1* gene expression is limited to a few specialized tissues, including brainstem raphe neurons, pinealocytes, the central nervous system (CNS), and part of the peripheral serotonergic nervous system [48]. Using a GWAS of 909 families (three members per family including ADHD patients and their parents), Sonuga-Barke et al. (2008) reported nominal evidence for interaction between *TPH1* rs623580 and parental criticism when predicting conduct disorder symptom [49]. Although the underlying mechanism was still unclear, this study provided the primary evidence for *TPH1* rs623580 might moderate the relation between adverse environments and outcomes. Therefore, the present study designed to employ *TPH1* rs623580 as the moderator to investigate whether it could moderate the relation between paternal indifference & neglect in early life and creativity in youth.

In summary, the current study aimed to explore the impact of paternal indifference & neglect in early life, *TPH1* rs623580, offspring’s gender, and the interaction effects thereof on creativity in youth. It is postulated that paternal indifference & neglect in early life would be negatively predict creativity in youth. It is also assumed that *TPH1* rs623580 polymorphism and offspring’s gender would moderate the negative influence of paternal indifference & neglect in early life on creativity in youth.

## METHODS

### Participants and Procedure

Participants included 539 (183 males and 356 females) unrelated healthy Han Chinese undergraduate students with an average age of 18.93 years (SD=1.084, range=17–22) from Shandong Normal University. None of the participants had been hospitalized for head trauma, psychiatric or neurologic reasons and none abused alcohol or drugs. The study protocol was approved by Institutional Review Board of Shandong Normal University. Written informed consent for genetic analysis was obtained from each participant after a description and explanation of the study.

### *TPH1* rs623580

DNA was extracted from peripheral venous blood samples using the QIAamp DNA Mini Kit (Qiagen, Valencia, CA, USA). Genotyping was carried out by a technician blind to other data from the research project. The single nucleotide polymorphisms (SNPs) were genotyped at the Beijing Genomics Institute-Shenzhen (BGI-Shenzhen, Shenzhen, China) using the Sequenom®MassARRAY®iPLEX system (Sequenom, San Diego, CA, USA). A customized set of SNPs was provided to BGI-Shenzhen by the investigator and BGI-Shenzhen provided the final oligonucleotides sequences to be used. Reverse and extension primers were designed using the MassARRAY Assay Design 3.0. For quality control, 5% random DNA samples were re-genotyped for each SNP, yielding a reproducibility of 100%. The *TPH1* rs623580 polymorphism was assessed as part of the SNP panel and met the criteria mentioned above. The genotype distribution of *TPH1* rs623580 for AA was 14.5% (n=78), AT was 50.2% (n=271), and TT was 35.3% (n=190). Consistent with previous research [50], AA and AT genotypes were combined and compared with the TT group. Allelic frequency of *TPH1* rs623580 is presented in Table 1.

**Table 1.**
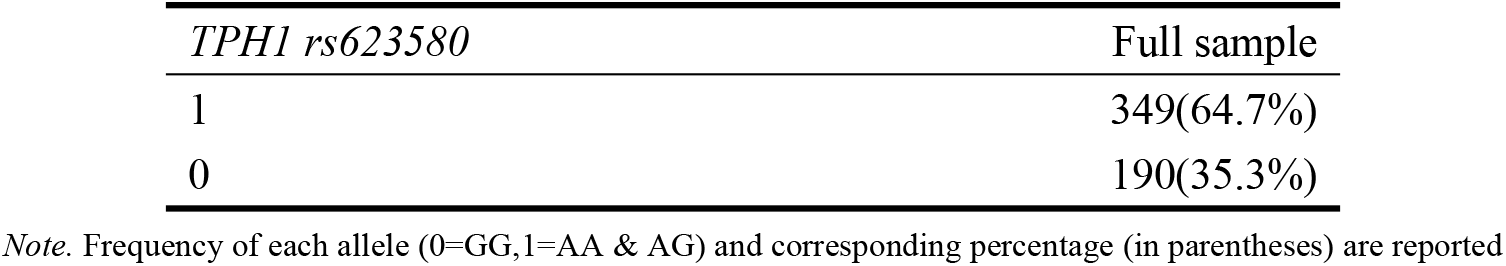
*Frequency of the TPH1 rs623580*

| *TPH1 rs623580* | Full sample |
| --- | --- |
| 1 | 349(64.7%) |
| 0 | 190(35.3%) |

*Note.* Frequency of each allele (0=GG,1=AA & AG) and corresponding percentage (in parentheses) are reported

## Measures

### Creative potential measures

Creativity was measured by Figural Divergent Thinking Test selected from the Runco Creativity Assessment Battery (rCAB; Creativity Testing Service, Bishop, GA, USA). Three line-drawings were represented in these tests, and participants were asked to list as many things as they can. Four minutes were allowed for each item. According to the guideline of Creativity Testing Service, the following three scores were obtained: fluency, flexibility, and originality. Fluency score was obtained by counting the number of unduplicated ideas provided by each participant. Originality score was calculated by counting the number of unusual ideas given by less the 5% of the sample. To score flexibility, a category list was first generated for each item, and the flexibility score was the number of different categories used in one participant’s ideas. Two trained raters (both were psychology graduate students from Shandong Normal University) were engaged to score all those ideas. The Chinese version of this measure was a widely used noninvasive measure and demonstrated adequate reliability and validity [3, 11, 20, 51, 52]. The inter-rater reliabilities for all the three scores in present study were higher than. 95; therefore, the final scores were obtained by averaging scores from the two raters. In current study, the Cronbach’s alpha was. 86 for fluency,. 69 for flexibility, and. 83 for originality.

### Parental Bonding Instrument (PBI)

The Parental Bonding Instrument is a 25-item self-rating questionnaire designed to measure the quality of the attachment or bond between parent and child, based on the memory of participants regarding their parents before their age of 16 [53]. Six items define the “care”, in which the higher the score, the higher the affection and warmth exercised by their parent; six items define the “indifference & neglect”, in which the higher the score, the higher the rejection and neglect exercised by their parent; seven items establish the “overprotection”, in which the higher the score, the higher the over involvement attitude and psychological control from parents; six items on the “autonomy”, in which the higher the score, the higher the encouragement of independence attitude and psychological autonomy from parents [54]. Participants scored each of their parents separately, on a 4-point Likert-type scale ranging from 0 (‘‘very unlike’’) to 3 (‘‘very like’’). The Chinese version of this measure was available and established reliability and validity [55]. In this study, care and indifference & neglect dimensions was used to measure the paternal rearing attitudes, the Cronbach’s alpha was. 84 for care, and. 78 for indifference & neglect.

### Data analysis

To test whether the relationships between paternal indifference & neglect and creativity (fluency, flexibility, originality) were moderated by *TPH1* rs623580 and offspring’s gender, a series of hierarchical regression analyses were performed. Paternal care was significantly related to paternal indifference & neglect and was therefore included in the regressions. Age and paternal care were included as covariates in the first regression step. In the second step, creativity (fluency, flexibility, originality) was predicted from the main effects of offspring’s gender (male coded as 1 and female as 0), paternal indifference & neglect, and *TPH1* rs623580. Then the moderator term (the interaction between paternal indifference & neglect, *TPH1* rs623580, and offspring’s gender) was added in the third step.

Because all three-way interaction effect on three outcome were not significant, we performed two two-way interaction separately on each outcome. When significant paternal indifference & neglect × *TPH1* rs623580 and *TPH1* rs623580 × offspring’s gender interactions were found, the nature of the interactions was tested by post-hoc analyses. The SPSS version 16.0 was used for analysis.

## Results

Table 2 reports the correlations, means, and standard deviations of the variables of this study. Paternal care were positively correlated with fluency (r=0.127, *p*<0.01), flexibility (r=0.112, *p*<0.01), and originality (r=0.117, *p*<0.01). Paternal indifference & neglect were negatively correlated with fluency (r=-0.107, *p*<0.05), flexibility (r=-0.085, *p*<0.05), and originality (r=-0.089, *p*<0.05). There were evidences for gender differences in fluency (r=-0.278, *p*<0.01), flexibility (r=-0.225, *p*<0.01), and originality (r=-0.195, *p*<0.01), but not in *TPH1* rs623580(r=-0.061, *p*>0.05) and each of those paternal bonding variables (*p*s>0.05). *TPH1* rs623580 was not correlated with any paternal bonding variables, offspring’s gender, and each of the outcome variables (*p*s>0.05). The findings of the interaction effect of paternal indifference & neglect and *TPH1* rs623580 on the outcome variables are summarized in Table 3. The findings of the interaction effect of *TPH1* rs623580 and offspring’s gender on the outcome variables are summarized in Table 4.

**Table 2.** *Correlations among primary study variables*

| Variable | 1 | 2 | 3 | 4 | 5 | 6 | 7 | 8 |
| --- | --- | --- | --- | --- | --- | --- | --- | --- |
| 1.age | — |  |  |  |  |  |  |  |
| 2.gender | .101* | — |  |  |  |  |  |  |
| 3.rs623580 | -.027 | -.061 | — |  |  |  |  |  |
| 4. PC | -.102* | -.045 | -.019 | (.84) |  |  |  |  |
| 5. PI | .077 | .075 | -.006 | -.741** | (.78) |  |  |  |
| 6. fluency | -.012 | -.278** | -.058 | .127** | -.107* | (.86) |  |  |
| 7. originality | -.004 | -.195** | -.050 | .117** | -.089* | .930** | (.83) |  |
| 8. flexibility | -.031 | -.225** | -.073 | .112** | -.085* | .819** | .741** | (.69) |
| Mean | 18.91 | .34 | .65 | 2.03 | .76 | 10.05 | 4.88 | 5.10 |
| SD | 1.08 | .47 | .48 | .59 | .54 | 4.20 | 3.00 | 1.26 |

*Note.* Male = 1, Female = 0; PC = Paternal care, PI = Paternal indifference & neglect; \* $p < .05$ , \*\* $p < .01$ .

**Table 3.** *Hierarchical linear regression analysis testing the effects of Paternal indifference & neglect, TPH genotype and their interaction on creativity*.

| Variables | Model 1<br>Fluency<br>$\beta$ | Model 2<br>Fluency<br>$\beta$ | Model 3<br>Fluency<br>$\beta$ | Model 4<br>Originality<br>$\beta$ | Model 5<br>Originality<br>$\beta$ | Model 6<br>Originality<br>$\beta$ | Model 7<br>Flexibility<br>$\beta$ | Model 8<br>Flexibility<br>$\beta$ | Model 9<br>Flexibility<br>$\beta$ |
| --- | --- | --- | --- | --- | --- | --- | --- | --- | --- |
| Age | .001 | .000 | -.010 | .008 | .006 | -.002 | -.020 | -.022 | -.031 |
| PC | .125** | .104 | .105 | .117** | .112 | .112 | .108* | .104 | .105 |
| PI |  | -.028 | .119 |  | -.005 | .132 |  | -.004 | .138 |
| rs623580 |  | -.056 | -.056 |  | -.047 | -.048 |  | -.071 | -.071 |
| PI × rs623580 | | | -.182* | | | -.170* | | | -.175* |
| F | 4.275* | 2.593* | 3.375** | 3.650* | 2.127 | 2.826* | 3.401* | 2.386* | 3.103** |
| R<sup>2</sup> | .016 | .019 | .031 | .013 | .016 | .026 | .013 | .018 | .028 |
| $\Delta R^2$ | .012 | .012 | .022 | .010 | .008 | .017 | .009 | .010 | .019 |

Note. Male = 1, Female = 0; PC= Paternal care, PI= Paternal indifference & neglect; \* $p < .05$ , \*\* $p < .01$ .

**Table 4.**
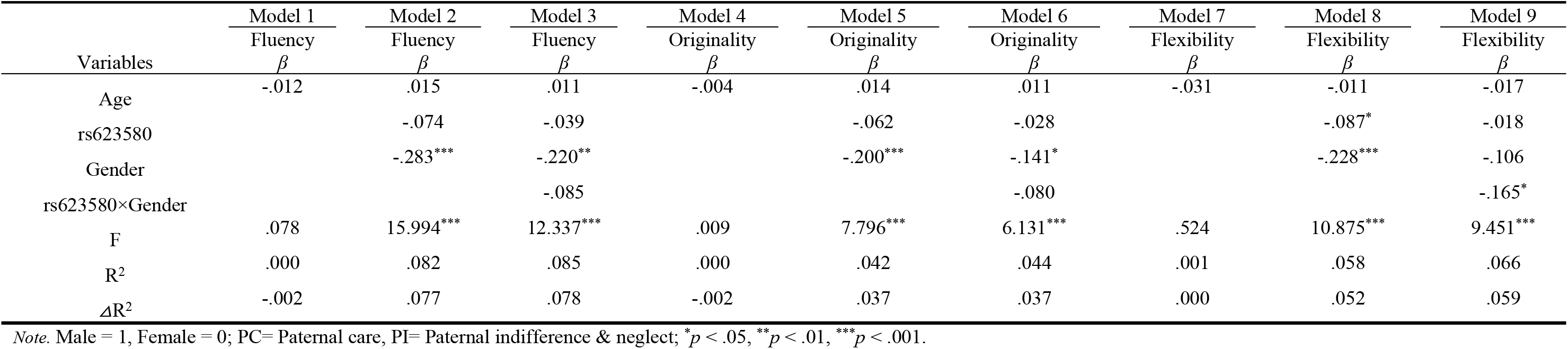
*Hierarchical linear regression analysis testing the effects of TPH genotype, gender and their interaction on creativity*.

| Variables | Model 1<br>Fluency<br>$\beta$ | Model 2<br>Fluency<br>$\beta$ | Model 3<br>Fluency<br>$\beta$ | Model 4<br>Originality<br>$\beta$ | Model 5<br>Originality<br>$\beta$ | Model 6<br>Originality<br>$\beta$ | Model 7<br>Flexibility<br>$\beta$ | Model 8<br>Flexibility<br>$\beta$ | Model 9<br>Flexibility<br>$\beta$ |
| --- | --- | --- | --- | --- | --- | --- | --- | --- | --- |
| Age | -.012 | .015 | .011 | -.004 | .014 | .011 | -.031 | -.011 | -.017 |
| rs623580 |  | -.074 | -.039 |  | -.062 | -.028 |  | -.087* | -.018 |
| Gender |  | -.283*** | -.220** |  | -.200*** | -.141* |  | -.228*** | -.106 |
| rs623580×Gender |  |  | -.085 |  |  | -.080 |  |  | -.165* |
| F | .078 | 15.994*** | 12.337*** | .009 | 7.796*** | 6.131*** | .524 | 10.875*** | 9.451*** |
| R<sup>2</sup> | .000 | .082 | .085 | .000 | .042 | .044 | .001 | .058 | .066 |
| $\Delta R^2$ | -.002 | .077 | .078 | -.002 | .037 | .037 | .000 | .052 | .059 |

Note. Male = 1, Female = 0; PC= Paternal care, PI= Paternal indifference & neglect; \* $p < .05$ , \*\* $p < .01$ , \*\*\* $p < .001$ .

### Paternal indifference & neglect and fluency: *TPH1* rs623580 and Offspring’s Gender as Moderators

Results showed that both paternal indifference & neglect and offspring’s gender had direct main effects on fluency (*B*=1.577*, p<0.05; B*=-1.936*, p<0.01*), and *TPH1* rs623580 did not had a direct main effect (AA & AT=1, *B*=-0.351, *p*=0.437). The three-way interaction of paternal indifference & neglect, offspring’s gender and *TPH1* rs623580 on fluency was not significant (*B*=0.371, *p*=0.788), but there was a significant two-way interaction of paternal indifference & neglect and *TPH1* rs623580 (*B*=-0.193, *p<0.05*). This two-way interaction remained significant after the non-significant three-way and all non-significant two-way interaction terms were dropped and a reduced model was run (*B*=-0.182, *p<0.05*) (see Table 3).

The significant interactions term of paternal indifference & neglect and *TPH1* rs623580 on fluency was tested for each *TPH1* genotype group. Results of the regression for AA / AT genotypes indicated that paternal indifference & neglect was related to lower level of fluency (*Β*=-1.429, *p*<0.05, 95% *CI*=-2.240 to −0.617). In contrast, results of the regression for TT genotype indicated that paternal indifference & neglect was not associated with fluency (*Β*=0.310, *p*>0.05, 95% *CI*=-0.787 to 1.407). Regression lines depicting levels of paternal indifference & neglect for AA / AT genotypes and TT genotype are plotted in Figure 1a.

**Figure 1.**
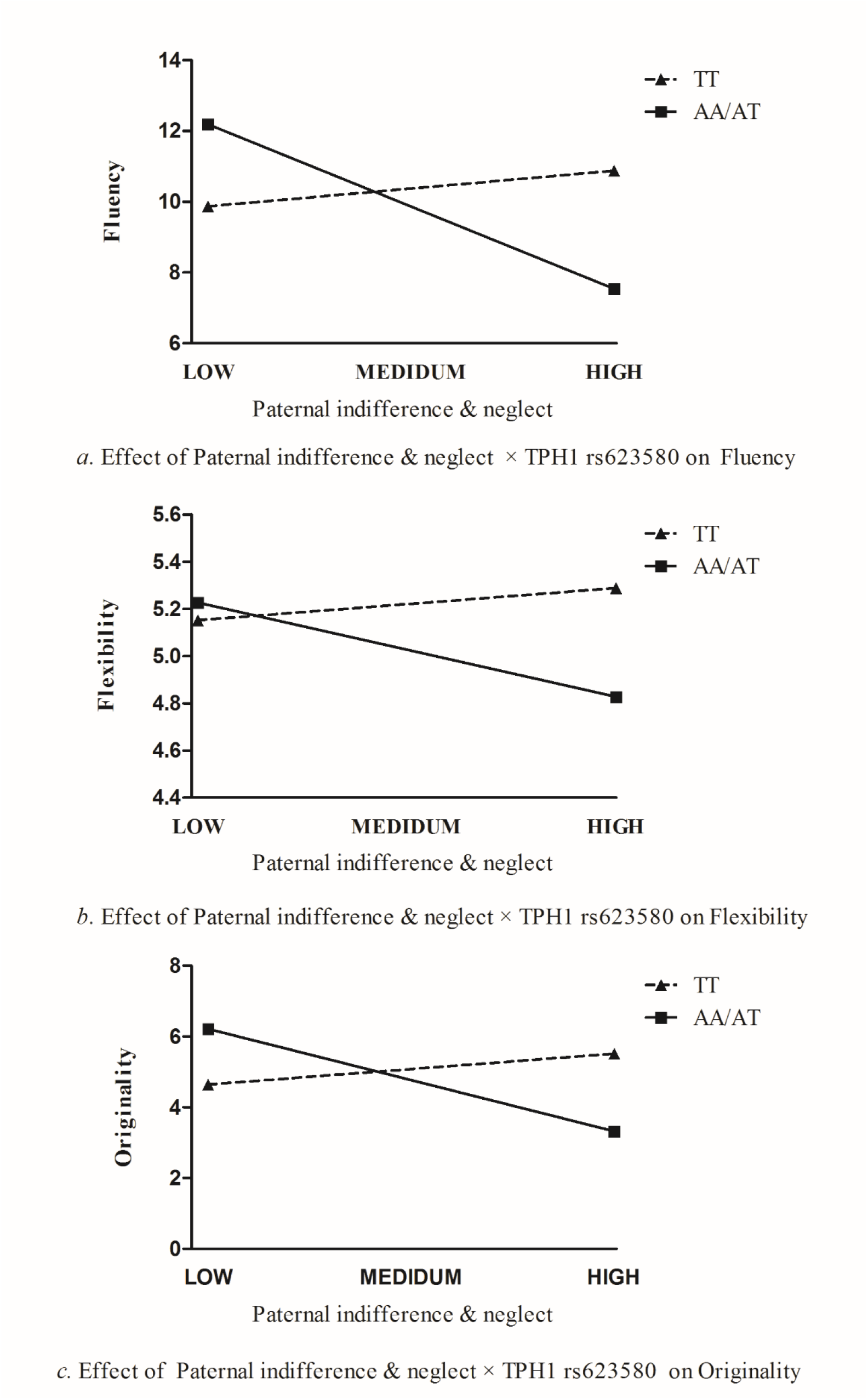
Effect of Paternal indifference× TPH1 rs623580 on Fluency, Flexibility, and Originality.

### Paternal indifference & neglect and originality: *TPH1* rs623580 and Offspring’s Gender as Moderators

Results showed that both paternal indifference & neglect and offspring’s gender had direct main effects on originality (*B*=1.253, *p*<0.05; *B*=-0.876, *p*<0.05), and the *TPH1* rs623580 did not had a direct main effect (AA & AT=1, *B*=-0.181, *p*=0.583). Although the three-way interaction of paternal indifference & neglect, offspring’s gender and *TPH1* rs623580 on originality was not significant (*B*=0.402, *p*=0.689), there was a significant two-way interaction of paternal indifference & neglect and *TPH1* rs623580 (*B*=-0.190, *p<0.05*). This two-way interaction remained significant after the non-significant three-way and all non-significant two-way interaction terms were dropped and a reduced model was run (*B*=-0.170, *p<0.05*) (see Table 3).

The significant interactions term of paternal indifference & neglect and *TPH1* rs623580 on originality was tested for each *TPH1* genotype group. Results of the regression for AA / AT genotypes indicated that paternal indifference & neglect was related to lower level of originality (*Β*=-0.892, *p*<0.05, 95% *CI*=-1.457 to −0.326). In contrast, results of the regression for TT genotype indicated that paternal indifference & neglect was not associated with originality (*Β*=0.269, *p*>0.05, 95% *CI*=-0.558 to 1.096). Regression lines depicting levels of paternal indifference & neglect for AA / AT genotypes and TT genotype are plotted in Figure 1b.

### Paternal indifference & neglect and flexibility: *TPH1* rs623580 and Offspring’s Gender as Moderators

Results revealed no significant main effects of paternal indifference & neglect (*B*=0.445, *p*=0.067), *TPH1* rs623580 (*B*=-0.050, *p*=0.718) and offspring’s gender (*B*=-0.283, *p*=0.124). The three-way interaction of paternal indifference & neglect, offspring’s gender and *TPH1* rs623580 on flexibility was not significant (*B*=0.205, *p*=0.625). However, two significant two-way interactions emerged.

First, there was a significant interaction of paternal indifference & neglect and *TPH1* rs623580 (*B*=-0.193, *p*<0.05). This two-way interaction remained significant after the non-significant three-way and all non-significant two-way interaction terms were dropped and a reduced model was run (*B*=-0.175, *p*<0.05) (see Table 3). The significant interactions term of paternal indifference & neglect and *TPH1* rs623580 on flexibility was tested for each *TPH1* genotype group. Results of the regression for AA / AT genotypes indicated that paternal indifference & neglect was related to lower level of flexibility (*Β*=-0.369, *p*<0.05, 95% *CI*=-0.610 to −0.128). In contrast, results of the regression for TT genotype indicated that paternal indifference & neglect was not associated with flexibility (*Β*=0.13, *p*>0.05, 95% *CI*=-0.211 to 0.464). Regression lines depicting levels of paternal indifference & neglect for AA / AT genotypes and TT genotype are plotted in Figure 1c.

Second, an interaction emerged between *TPH1* rs623580 and offspring’s gender (*B*=-0.159, *p*<0.05). This two-way interaction remained significant after the non-significant three-way and all non-significant two-way interaction terms were dropped and a reduced model was run (*B*=-0.165, *p*<0.05) (see Table 4). The significant interactions term of *TPH1* rs623580 and offspring’s gender on flexibility was tested for each *TPH1* genotype group. Results of the regression for AA / AT genotypes indicated that male was related to lower level of flexibility (*Β*=-0.801 *p*<0.001, 95% *CI*=-1.073 to −0.529). In contrast, results of the regression for TT genotype indicated that offspring’s gender was not associated with flexibility (*Β*=-0.291, *p*>0.05, 95% *CI*=-0.660 to 0.078). Regression lines depicting levels of offspring’s gender for AA / AT genotypes and TT genotype are plotted in Figure 2.

**Figure 2.**
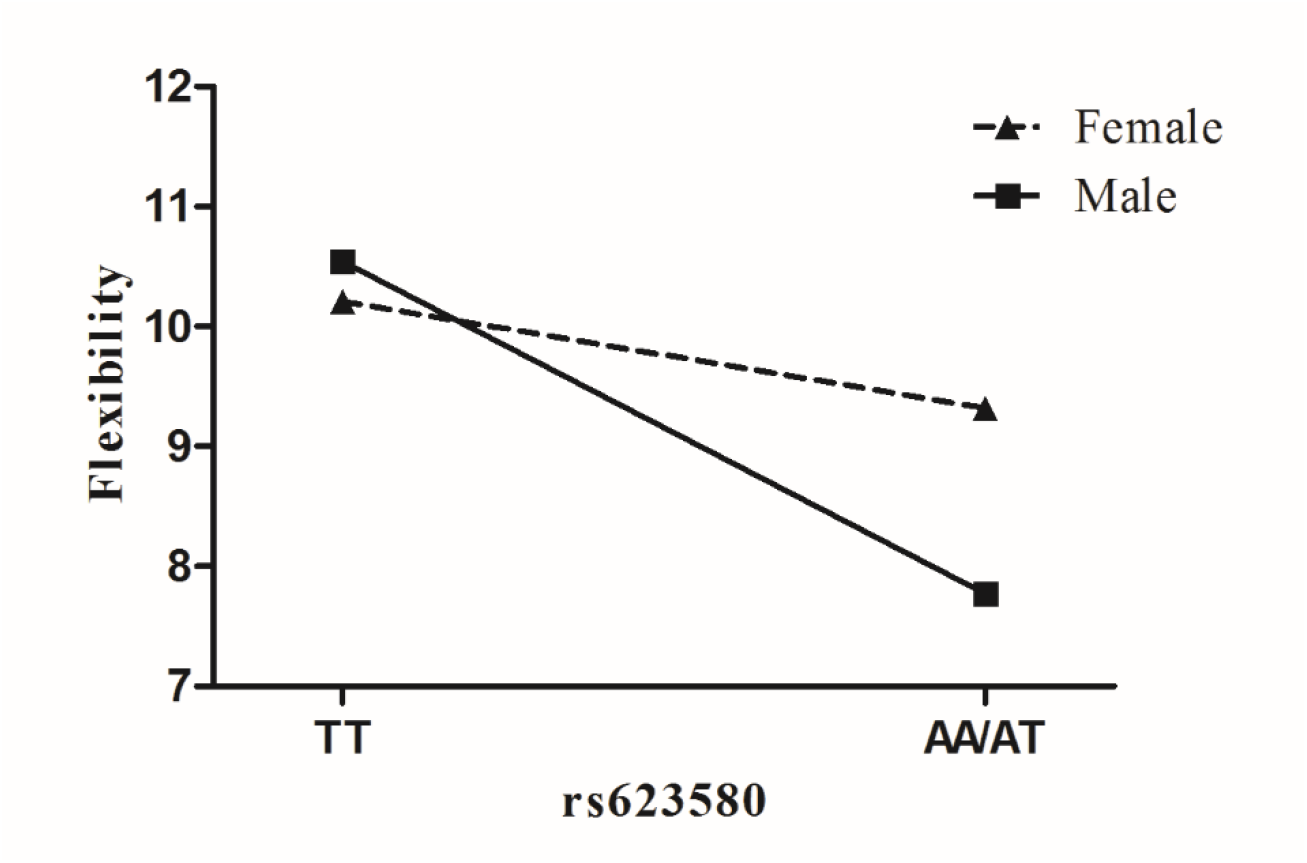
Effect of TPH1 rs623580 × Gender on Flexibility.

## Discussion

This study sought to examine the impact of paternal indifference & neglect in early life, *TPH1* rs623580, offspring’s gender, and the interaction effects thereof on creativity in youth. Two primary findings emerged. First, paternal indifference & neglect in early life negatively predicted creativity (fluency, flexibility and originality) in youth when individuals carry A allele of *TPH1* rs623580. Second, male offspring who carry A allele of *TPH1* rs623580 were linked with lower level of flexibility compared to TT homozygote individuals.

Firstly, present study provided supporting evidence for paternal indifference & neglect in early life negatively predicted on creativity (fluency and originality) in youth. These findings are consistent with previous research with Chinese samples which demonstrated that paternal rejection was negatively associated with adolescents’ creativity [56]. Given that indifferent and neglecting father usually remains psychologically and physically unresponsive or even inaccessible, they may be prejudicial to child’s psychological security [25]. Psychological security has been demonstrated positively predicted creativity [57, 58]. Therefore, it is reasonable to speculate that paternal indifference & neglect in early life may be adverse to individual’s psychological security, which has negative impact on creativity in youth. These findings of the direct effects of paternal indifference & neglect in early life on creativity in youth were congruent with prior studies in Western settings [34, 35].

Considering Chinese culture is widely characterized as collectivistic which emphasize interpersonal relatedness in contrast with Westernized cultures [59, 60]. Children might be more sensitive to paternal indifference/neglect, and perceive be rejected and lacking of paternal involvement and support in Chinese societies than in Western societies [25, 61]. However, it was difficult to compare the correlations for the two cultural groups due to lack of data on the correlations between paternal indifference & neglect and creativity in Western studies. Further examination of this issue is needed in future cross-cultural research.

Second, consistent with our expectation, a paternal indifference & neglect × *TPH1* rs623580 interaction was observed. It was found that the negative influence of paternal indifference & neglect in early life on creativity in youth was only present in individuals who carry A allele of *TPH1* rs623580 but not the carriers of the TT genotype, suggesting a hypothesis that carrying the A allele of *TPH1* rs623580 may increase the vulnerability to the early life adverse environments, paternal indifference & neglect for example, and pose a risk for creativity in youth. Paternal indifference & neglect in early life is identified as potent sources of stress, and is suggested to have a pervasive influence on a child’s psychological and biological regulatory processes [62]. Molecular genetics research demonstrated that *TPH1* mRNA expresses in the hypothalamus and the neuronal *TPH1* protein expresses in the anterior pituitary, these findings suggested that *TPH1* might involve in hypothalamic-pituitary-adrenal axis (HPA) regulation and influence on neuronal mechanisms of the brain [63, 64], including stress-response mechanisms [65]. Although *TPH1* rs623580 does not result in an amino acid substitution as located in a regulatory region, it may affects in *TPH1* enzyme activity [48]. Previous study have reported that *TPH1* rs623580 related to major depressive disorder (MDD) [66]. Most recent study further demonstrated that the A allele of *TPH1* rs623580 might increase the risk of depressive disorder [67].

Therefore, it is possible that *TPH1* rs623580 may moderate the negative relation between paternal indifference & neglect in early life and creativity in youth via regulating the stress-response processes. Specifically, compared with the TT homozygote individuals, the A allele carriers may have less capacity to withstand the corrosive drizzle of paternal indifference & neglect in early life and to cope with stress effectively, which in turn lead them to the damaging consequences [68, 69].

Third, a *TPH1* rs623580 × offspring’s gender interaction predicting flexibility was detected in the present study. Specifically, males who carrying the A allele showed lower flexibility than the TT carriers. This result further supported the hypothesis that A allele of *TPH1* rs623580 might be a risk allele for creativity, at least in males. Animal research indicated that sex hormones, including estrogen and progesterone, can increase *TPH1* expression in the central nervous system of primates [70]. It could be speculated that the significant effect of *TPH1* rs623580 A allele in male in the present study might due to poor sex hormones regulation because of lower level of estrogen and progesterone in male. Although the underlying mechanism of the interaction effect is not yet clear, the result suggested that *TPH1* rs623580 might involve in gender difference in creativity.

Several limitations of this study should be addressed. Firstly, the present study employed a retrospective design to explore the negative influence of paternal indifference & neglect in early life on creativity. Longitudinal study from early childhood to young adulthood was needed to understand the dynamic association between early life family environment and creativity. Secondly, the assessment of early life parental indifference & neglect in present study was limited in self-report measure, which might only reflect participants’ perceived parental indifference & neglect, not objectively observed parental indifference & neglect. Future study simultaneously including the parents and observer reports of early life family environment would provide more convincing results. Third, the present study used a relatively homogenous sample consisting of Chinese undergraduate students. As the genetic backgrounds vary for different ethnic populations, the generalization of the present findings to other samples is limited. Future research across populations of different genetic and cultural backgrounds are warranted to examine what extent the present findings can be generalized to other samples.

These limitations notwithstanding, some valuable information can be derived from our findings. Drawing upon gene × environment and gene × gender interaction research, this study provided evidence that carrying A allele of *TPH1* rs623580 might be a significant risk factor of creativity. The findings of the present study contribute to further understanding the role of genetic factors in the pathways that how the early life family environment shapes creativity in adulthood. In addition, our findings may also provide a new perspective to reevaluate the genetic basis of gender difference in creativity.

## Notes

Author Note Qi Yu, Si Si, Shun Zhang, and Jinghuan Zhang, Department of Psychology, Shandong Normal University, Jinan, China. This research was supported by National Natural Science Foundation of China (31470999, 31771235), Key special project of national key research and development program of China (SQ2017YFB1400102), Shandong Provincial Institute of Qilu Cultural Studies. We appreciate Dr. Mark A. Runco (Torrance Creativity Center, University of Georgia) for the directions and help about the Divergent Thinking Test scoring.

